# Elevating the status of code in ecology

**DOI:** 10.1101/027110

**Authors:** K. A. S. Mislan, Jeffrey M. Heer, Ethan P. White

**Author notes:** Corresponding author Email address (K. A. S. Mislan).

## Introduction

Code is increasingly central to research in ecology. From running statistical analyses for field and lab projects to developing software intended to be used broadly for modeling and data analysis, most ecologists now commonly write code as part of their research. It is important that the communication of research results addresses the fact that code is now a key component of most ecological studies.

## The role of code in ecological research

The transition to a greater reliance on code has been driven by increases in the quantity and types of data used in ecological studies, alongside improvements in computing power and software [1]. Code is written in programming languages like R and Python and is used by ecologists for a wide variety of tasks including manipulating, analyzing, and graphing data. A benefit of this transition to code-based analyses is that code provides a precise record of what has been done, making it easy to reproduce, adapt, and expand existing analyses.

Scientific code can be separated into two general categories - analysis code and scientific software. Analysis code is code that is used to correct errors in data, simulate model results, conduct statistical analyses, and create figures [2]. Release of analysis code is necessary for the results of a study to be reproducible. The majority of code written for ecological studies will likely be classified as analysis code and making this type of code available is extremely valuable even if it has not been refined to the highest standards [2, 3]. Scientific software is more general and is designed to be used in many different projects (e.g., R and Python packages). The development of ecological software is becoming more common and software is increasingly recognized as a research product [4, 5].

## Current standards for code in ecology

Journals are the primary method that ecologists use to communicate results of studies. Therefore, the way journals handle code is of import for evaluating the current status of code in ecology. To explore the current status of code in ecology journals, we identified journals through a search of the Journal Citation Reports (JCR) using the following search terms: “Ecology” for category, “2013” for year, “SCIE” (Science Citation Index) and “SSCI” (Social Sciences Citation Index) editions checked, and “Web of Science” for the category schema. We selected the top 100 results for analysis and, after excluding museum bulletins, a book, and a journal with broken website links, evaluated a total of 96 journals. We searched the author guidelines for each journal to determine if there was any mention of code or software in the context of scientific research. We also conducted more specific searches to determine if journals had a section for documentation of scientific software releases, and if journals had a policy requiring the release of code and/or data for article publication. Policies for the release of data are interesting to compare to the policies for the release of code because there has been an on-going community push for scientists to release data once results are published.

As of June 1, 2015, more than 75% of ecology journals do not mention scientific code in the author guidelines (Figure 1). Of the journals that mention scientific code, 14% require code to be made available. Nearly three times as many journals (38%) require data to be made available. A very small subset of journals (7%) have created a special section for software releases or added software releases to a list of options for existing methods sections (Figure 1). These findings are similar to recent studies of journal code policies in other scientific fields [6].

**Figure 1:**
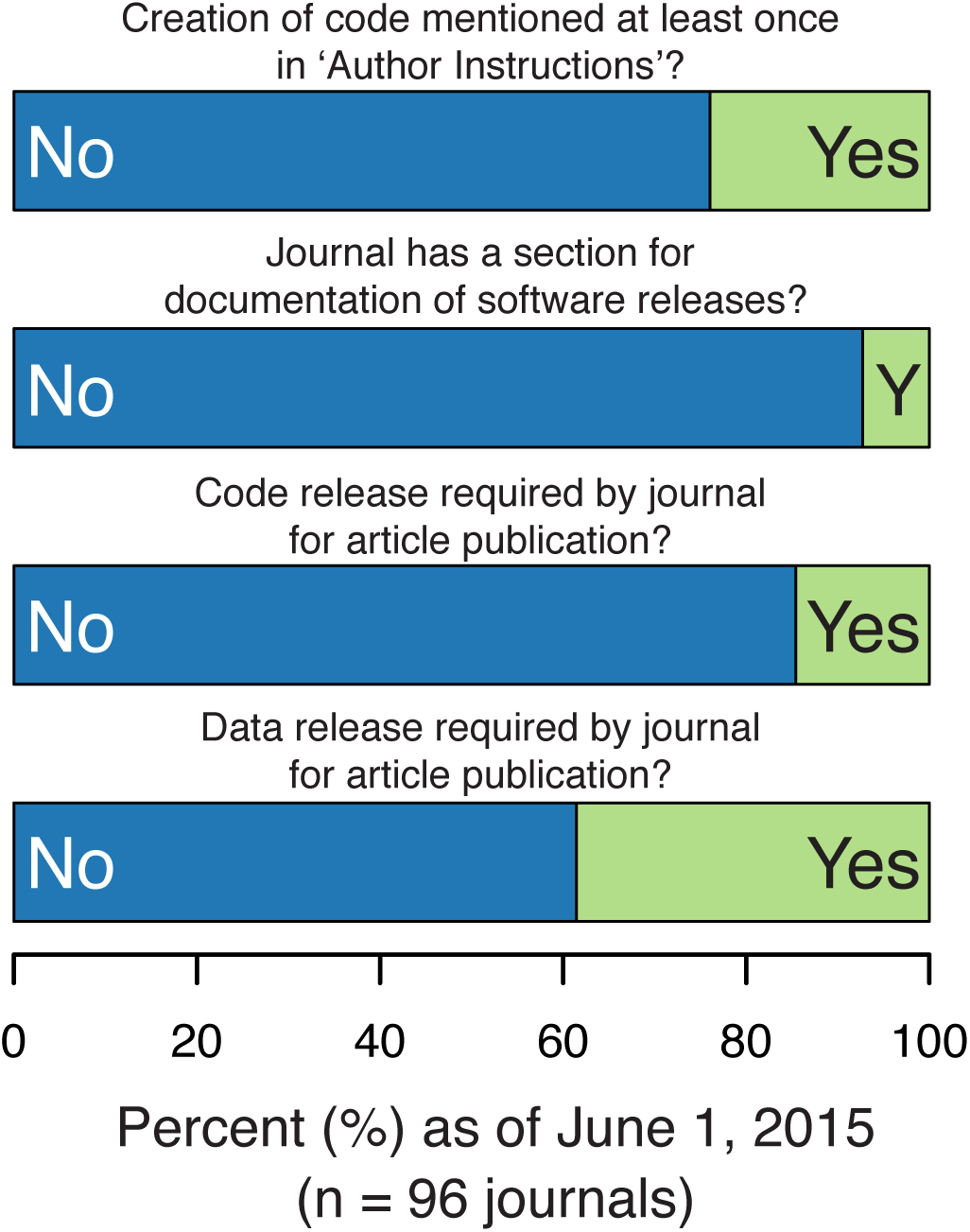
Most ecology journals do not have requirements or guidelines (as of June 1, 2015) for making code and data available. Ecology journals listed in the Journal Citation Reports (JCR) in 2013 were evaluated.

## Promoting code in ecology journals

Journals can promote the release of code used in ecological studies though a combination of increasing the visibility and discoverability of code and software and requiring code archiving. One way to increase visibility is to indicate code availability in the table of contents of all formats of the journal and have direct links from the online table of contents to the code (Figure 2a). In the article, links to code prominently displayed on the first page will also increase visibility (Figure 2b). This article format has already been adopted by some ecological journals for data, including *The American Naturalist.* In addition, journals can require and verify that code is made available at the time an article is submitted for review or is accepted for publication [7]. Requirements by journals for data to be made available have been very successful [3]. Specialized software sections in journals go a step further in promoting highly refined code that can be used broadly for ecological analyses and visualization [8]. Communicating the availability of software in a well-described journal format to the ecological community highlights software as a product of ecological research. Discoverability can be enhanced by having searchable databases for articles (e.g. journal archives, Web of Science, and PubMed) which include an option for selecting for articles with code. This search capability would make it much more feasible for an ecologist to find, compare, and adapt code from multiple research articles for a new study.

**Figure 2:**
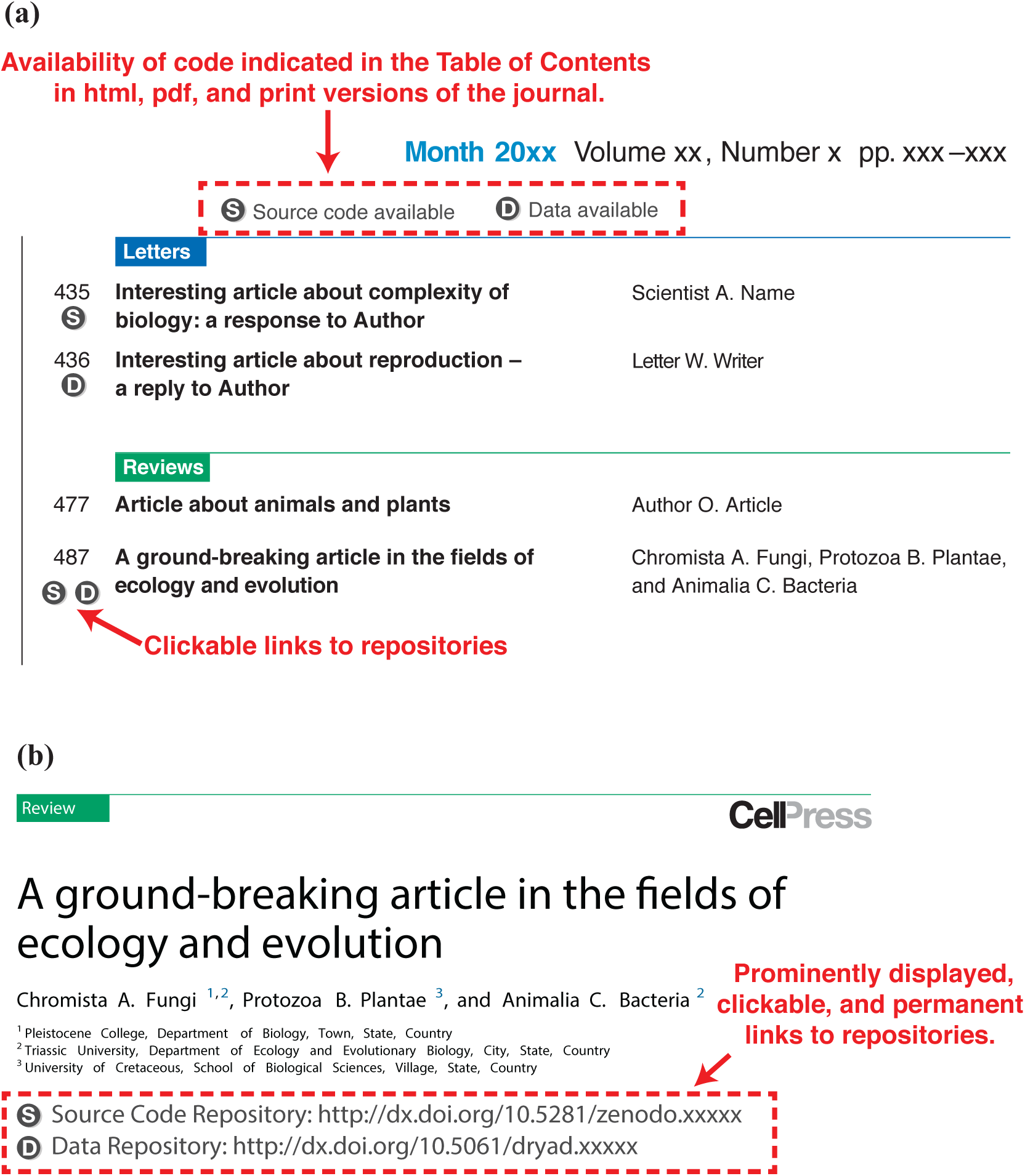
Recommended page layouts for: a) the table of contents of a journal; b) the first page of a journal article. The recommendations are for all formats of the journal including html, pdf, and print versions. An important feature is that active links can be clicked in electronic versions to directly access the code. The article titles and author names were made-up for the examples.

Ecologists may not be aware of the steps needed to share code or the ease of doing so with available resources [3]. Recent articles provide detailed information on the best practices for coding in the sciences and serve as essential guides for writing better code and sharing it with others [9, 10]. The incorporation of code into the review structure for articles is still an open question, but, at minimum, tests of the functionality of scientific code should be completed by the authors [5]. A critical step for sharing code is to put code in an archive that is open source, long-term, and citable, which will help ensure that code is widely available [5, 11]. Archiving options include code-specific repositories, data repositories, or the supplementary material of the journal itself.

Journals can have a significant impact on increasing the value of code within the ecological community. We believe that broad adoption of the suggestions to increase visibility and discoverability of code, as well as requiring its archiving, will motivate more authors to share code. By fostering reproducibility and reuse, more available code can improve the quality and accelerate the rate of research in ecology.

## Acknowledgements

K.A.S. was supported by the Washington Research Foundation Fund for Innovation in Data-Intensive Discovery and the Moore/Sloan Data Science Environments Project at the University of Washington. This work was partially supported by the Gordon and Betty Moore Foundation’s Data-Driven Discovery Initiative through grant GBMF4553 to J.M.H. and grant GBMF4563 to E.P.W.

